# Anti-seizure Gene Therapy for Focal Cortical Dysplasia

**DOI:** 10.1101/2023.01.09.523292

**Authors:** Amanda Almacellas Barbanoj, Robert T. Graham, Benito Maffei, Jenna C. Carpenter, Marco Leite, Justin Hoke, Felisia Hardjo, James Scott-Solache, Christos Chimonides, Stephanie Schorge, Dimitri M. Kullmann, Vincent Magloire, Gabriele Lignani

**Author notes:** Correspondence to: Gabriele Lignani, Vincent Magloire, Dimitri Kullmann.

## Abstract

Focal Cortical Dysplasias (FCDs) are a common subtype of malformation of cortical development, which frequently present with a spectrum of cognitive and behavioural abnormalities as well as pharmacoresistant epilepsy. FCD type II is typically caused by somatic mutations resulting in mTOR hyperactivity, and is the commonest pathology found in children undergoing epilepsy surgery. However, surgical resection does not always result in seizure freedom, and is often precluded by proximity to eloquent brain regions. Gene therapy is a promising potential alternative treatment and may be appropriate in cases that represent an unacceptable surgical risk. Here, we evaluated a gene therapy based on overexpression of the Kv1.1 potassium channel in a mouse model of frontal lobe FCD. An engineered potassium channel (EKC) transgene was placed under control of a human promoter that biases expression towards principal neurons (*CAMK2A*) and packaged in an adeno-associated viral vector (AAV9). We used an established FCD model generated by *in utero* electroporation of frontal lobe neural progenitors with a constitutively active human *RHEB* plasmid, an activator of mTOR Complex 1. First, we further characterised this by quantifying electrocorticograms and behavioural abnormalities, both in mice developing spontaneous generalised seizures and in mice only exhibiting abnormal interictal discharges. Then, using continuous video-electrocorticogram recordings from epileptic mice before and after injection of AAV9-*CAMK2A*-EKC in the dysplastic region, we observed a robust decrease in the frequency of seizures and in interictal activity, compared to mice injected with a control viral vector. Despite the robust anti-epileptic effect of the treatment, there was neither an improvement nor a worsening of performance in behavioural tests sensitive to frontal lobe function. AAV9-*CAMK2A*-EKC had no effect on interictal activity or behaviour in non-epileptic mice. AAV9-*CAMK2A*-EKC gene therapy is a promising therapy with translational potential to treat the epileptic phenotype of mTOR-related malformations of cortical development. Cognitive and behavioural co-morbidities may, however, resist an intervention aimed at reducing circuit excitability.

## Introduction

Focal Cortical Dysplasia (FCD) is a malformation of cortical development (MCD) and is among the leading causes of drug-resistant focal epilepsy^1^ pathology found in children undergoing epilepsy surgery^2^. Because tissue is required to confirm the diagnosis, precise estimates of the prevalence are not available. Nevertheless, the European Epilepsy Brain Bank reported that 20% of specimens collected from therapeutic surgery corresponded to an MCD, of which 71% were classified as FCD^1^.

Surgical resection of the dysplastic focus is often indicated to treat patients with pharmacoresistant epilepsy associated with FCD. However, although surgery can be successful in 14-63% of operated cases^3^, it is contraindicated in the majority of patients because of risks to normal brain function. There is therefore an urgent need for an effective therapy for FCD-associated epilepsy. The present study set out to test a gene therapy in a preclinical model.

FCD is classified histologically into several subtypes. FCD type II (FCD II) is characterized by dyslamination and dysmorphic neurons in a restricted area of the cortex and is generally caused by somatic mutations in neural progenitors^4^. Approximately 60% of FCD II lesions are caused by mutations affecting the mTORC1 (mammalian Target of Rapamycin Complex 1) signalling pathway^1^. Despite advances in the molecular defects underlying FCD II, the mechanisms leading to seizures are incompletely understood^1^, as are the circuit disorders underlying cognitive and behavioural comorbidities that are frequently seen in affected children.

In recent years, new animal models of FCDs have been developed, which rely on activating the mTORC1 pathway through either the downregulation of mTORC1 inhibitors (DEPDC5^5^, PTEN^6^, TSC1/2^7^) or upregulation of mTORC1 activators (Akt^8^, Rheb^9,10^, mTOR^11^). The tools used to modulate this pathway include CRISPR-Cas genome editing, cell-specific promoters, and *in-utero* electroporation. These techniques allow for a focal, mosaic dysregulation of cortical development, giving these animal models construct and face validity.

In this study, we used a mouse model of mTORC1-related FCD II generated by *in-utero* electroporation of neuronal progenitors with a plasmid carrying a Ras homolog mTORC1 binding (*RHEB*) gene with a mutation (S16H) that increases mTORC1 activity^9^. The *RHEB^CA^* (constitutively active *RHEB*) plasmid was targeted to the frontal cortex unilaterally to mimic FCD II.

Overexpression of the voltage gated potassium channel Kv1.1, encoded by *KCNA1*, leads to a moderate decrease in neuronal excitability and neurotransmitter release from axonal terminals^12^. Consequently, it has been validated as an effective gene therapy in several rodent models of focal epilepsy (focal neocortical epilepsy^13^ and temporal lobe epilepsy^14,15^), without off-target effects on a range of behaviours. Robust reductions in seizure frequency and/or duration were achieved by overexpressing wild type *KCNA1* or an engineered version that bypasses the normal post-transcriptional editing of *KCNA1* mRNA and reduces inactivation (engineered potassium channel, or EKC) ^14^. An anti-epileptic effect has also been demonstrated by CRISPR-mediated upregulation of the endogenous murine *Kcna1* gene^16^, and recently by using activity-dependent promoters to drive EKC^15^.

Given the robust evidence for anti-seizure effects of exogenous EKC delivery in other models, we asked if it has a similar potential therapeutic effect in FCD II, and whether it affects cognitive and behavioural co-morbidities. We observed a substantial reduction in spontaneous seizures with no alteration in behavioural tests sensitive to frontal lobe function. EKC overexpression is thus a promising gene therapy to reduce seizure burden in FCDs. The absence of effect on behavioural tests supports the safety of the treatment, while underlining that co-morbidities may reflect a broader disorder of brain function in FCD.

## Results

### The *RHEB*^CA^ mouse model recapitulates histological, electrophysiological and cognitive hallmarks of FCD II

We first refined a previously reported model of FCD^9^. We used a 3-electrode *in utero* electroporation design^17^ with a plasmid expressing *RHEB*^CA^ together with a tandem dimer Tomato (tdTomato) fluorescent reporter linked by a cleavable T2A peptide (pCAG-tdTomato-T2A-*RHEB*^CA^). Mouse embryos were electroporated at E14 ± 0.5 and pups were screened with transcranial epifluorescence imaging at P21 to confirm unilateral tdTomato expression in the frontal lobe (**Fig. 1A**). As previously reported in this mouse model^9^, we observed several characteristic features of FCD II in a subset of animals used for histology; electroporated neurons were heterotopic, dysmorphic, and cytomegalic **(Fig. 1B-E)**. Indeed, the average soma size of *RHEB*^CA^ cells was almost three-fold greater than neurons electroporated with a control plasmid expressing only tdTomato **(Fig. 1B)**. We also observed migration defects **(Fig. 1C)** and an increase in mTORC1 activity assessed by the quantification of S6 protein phosphorylation compared to the contralateral hemisphere **(Fig. 1D, E)**.

**Figure 1.**
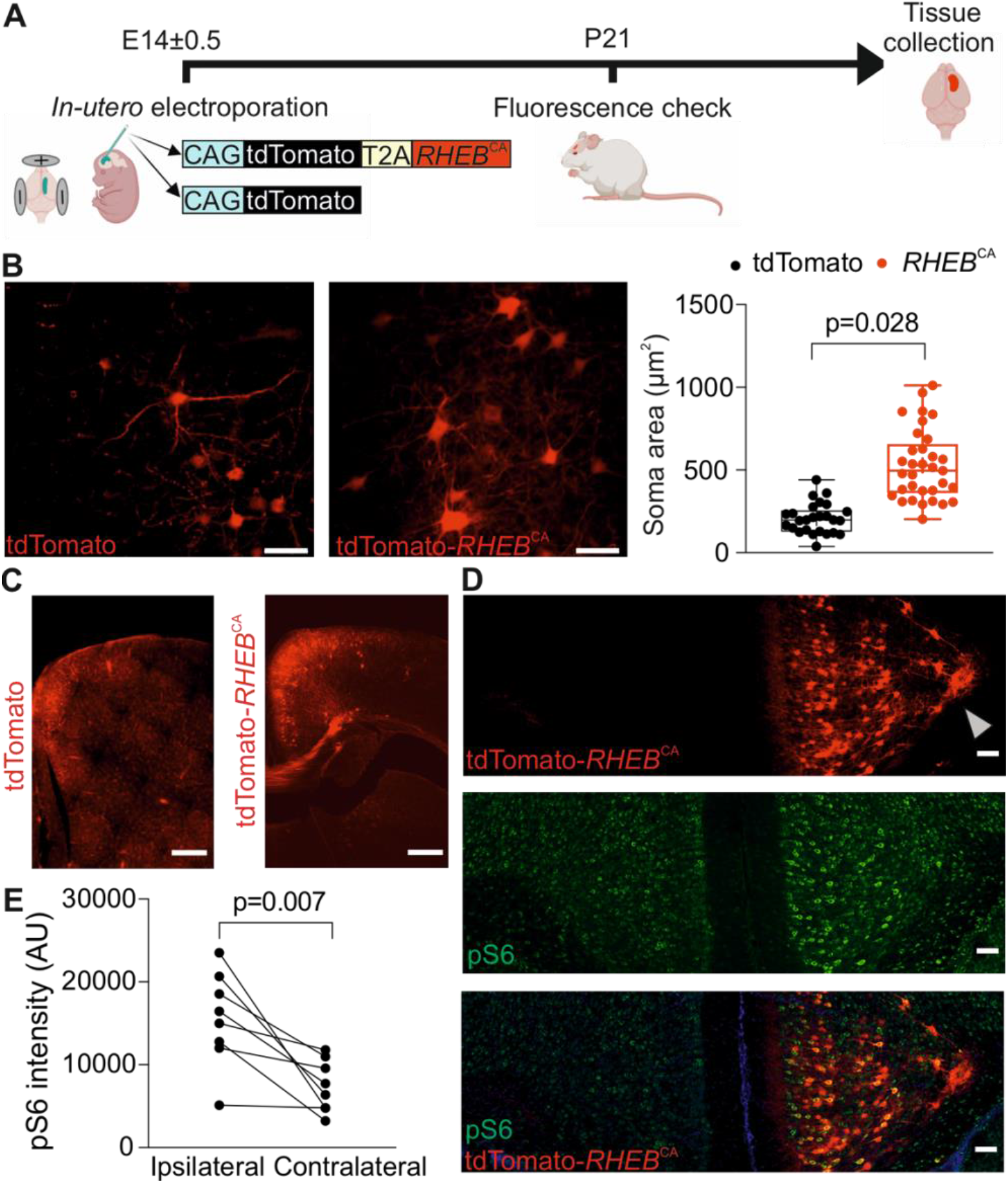
*RHEB*^CA^ mouse model recapitulates histological hallmarks of FCD II. **A.** Experimental Plan. **B.** (*left and centre panels*) Fluorescence images of neurons expressing either tdTomato alone (control) or tdTomato-*RHEB*^CA^, scale bars = 50μm. (*right panel*) Quantification of neuronal soma area (tdTomato, n=26 cells from 2 mice; *RHEB*^CA^, n = 33 cells from 2 animals, unpaired Student’s t-test). **C.** tdTomato or tdTomato-*RHEB*^CA^ prefrontal cortical slices, scale bars = 500μm. **D.** Representative coronal sections of prefrontal cortex in a *RHEB* electroporated mouse. *Top:* tdTomato; *middle:* pS6; *bottom:* merged images. Note the dyslamination and heterotopic neurons on the top panel (arrowhead: heterotopic neurons in the corpus callosum), scale bars = 100 μm. **E.** pS6 fluorescence intensity in the ipsilateral (electroporated) and contralateral hemispheres (n = 8 slices from 2 animals, paired Student’s t-test).

We next characterised the phenotype of *RHEB*^CA^ electroporated animals. For this, we implanted subcutaneous wireless electrocorticogram (ECoG) transmitters with a recording electrode over the tdTomato-positive area. After 15 days of baseline recording, animals underwent behavioural tests **(Fig. 2A).** A subset of animals underwent concurrent video monitoring, which showed that seizures detected using a custom algorithm corresponded to generalized tonic-clonic seizures with convulsions (Racine scale 5). These generalised seizures started with behavioural arrest, followed by either forelimb jerking or stereotypical rotation before generalised tonic-clonic activity with loss of postural tone (supplementary video 1).

**Figure 2.**
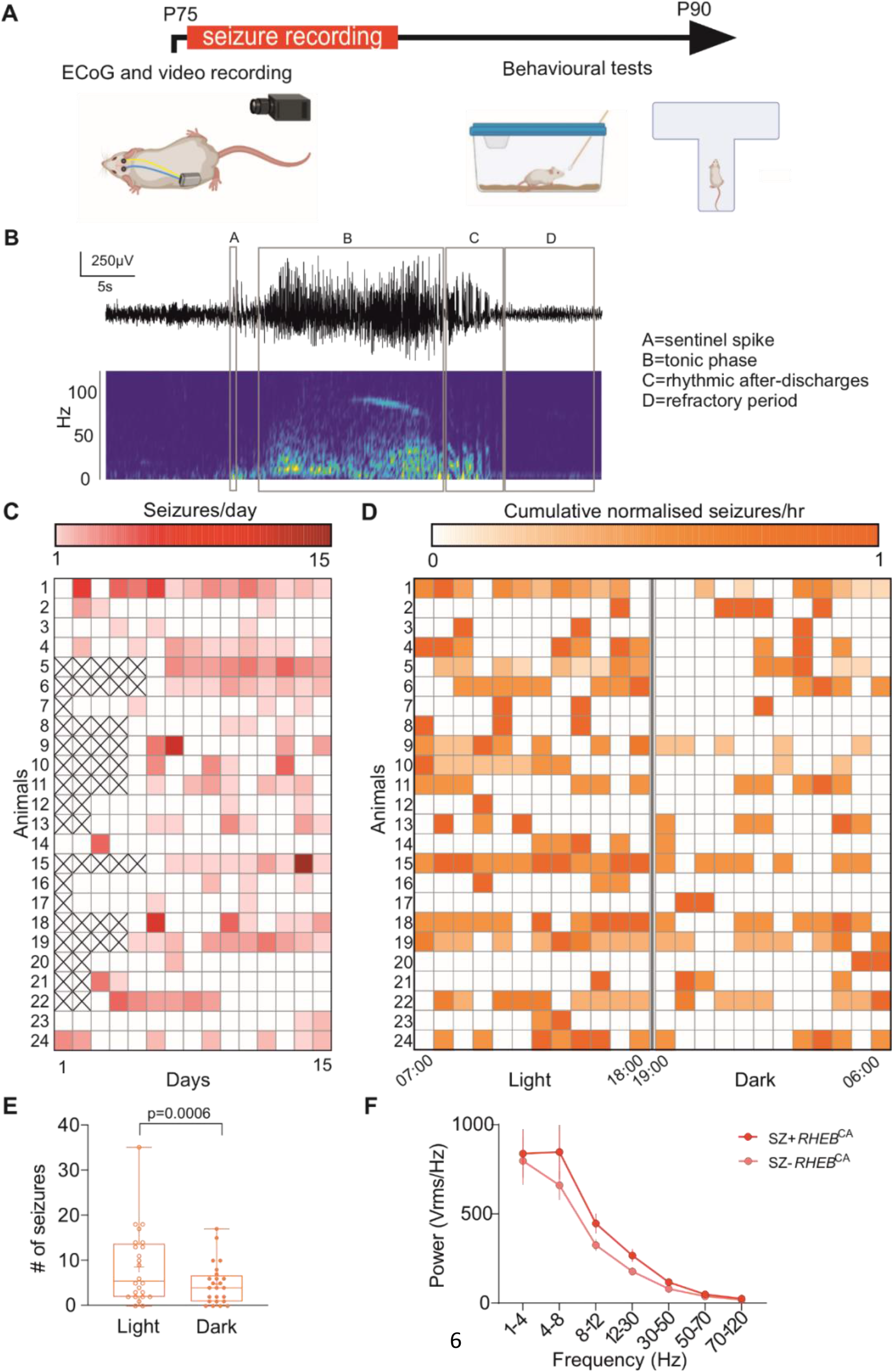
*RHEB*^CA^ electroporated mice exhibit spontaneous generalised seizures preferentially during the light phase of the 24hr light-dark cycle. **A.** Timeline showing ECoG and video recording for quantification of seizures and behavioural tests. **B.** Representative seizure displaying the 4 main features used for automatic detection. **C.** Raster plot showing the number for seizures for each animal (15 days, n=24). **D.** Raster plot of the distribution of the number of seizures per hour for each animal during the entire baseline recording period, normalised for each animal to the day with the maximum number of seizures (15 days, n=24 animals). **E.** Box plot showing seizure occurrence during the light and dark periods (n=24, paired Student’s t-test). **F.** Power at different frequency bands in animals with (SZ+ *RHEB*^CA^ n=24) and without (SZ- *RHEB*^CA^ n=13) seizures.

Overall, we observed generalised seizures in 24 out of 37 *RHEB^CA^*-electroporated animals (~65%). Epileptic animals had a median frequency of 0.9 seizure per day (**Fig. 2B, C**), less than previously reported using a similar experimental design^18^. Among possible explanations for the lower seizure frequency is the introduction of the T2A linker in the plasmid construct^19^. Epileptic animals exhibited ~57% seizure-free days during a 15-day recording period (**Fig. 2C**). Seizure occurrence was greatest during the light phase of the 12h:12h light-dark cycle (**Fig. 2D, E**). Given that mice tend to be more active during the dark phase, this finding is reminiscent of the propensity for nocturnal seizures in patients with frontal lobe epilepsy^20^. We also observed a trend towards an increase in ECoG power in animals with seizures compared to those without (SZ+ *RHEB*^CA^ and SZ- *RHEB*^CA^, respectively) (**Fig. 2F**).

We also examined the occurrence of interictal spikes (**Fig. 3A**), which are a frequent biomarker of focal epilepsy^21^, in *RHEB*^CA^ electroporated animals either with (SZ+ *RHEB*^CA^) or without (SZ- *RHEB*^CA^) generalised seizures. In contrast to seizures, interictal spikes displayed no obvious relationship to the light-dark cycle (**Fig. 3B**). However, SZ+ *RHEB*^CA^ animals had a higher frequency of spikes than SZ- *RHEB*^CA^ mice (**Fig. 3C**).

**Figure 3.**
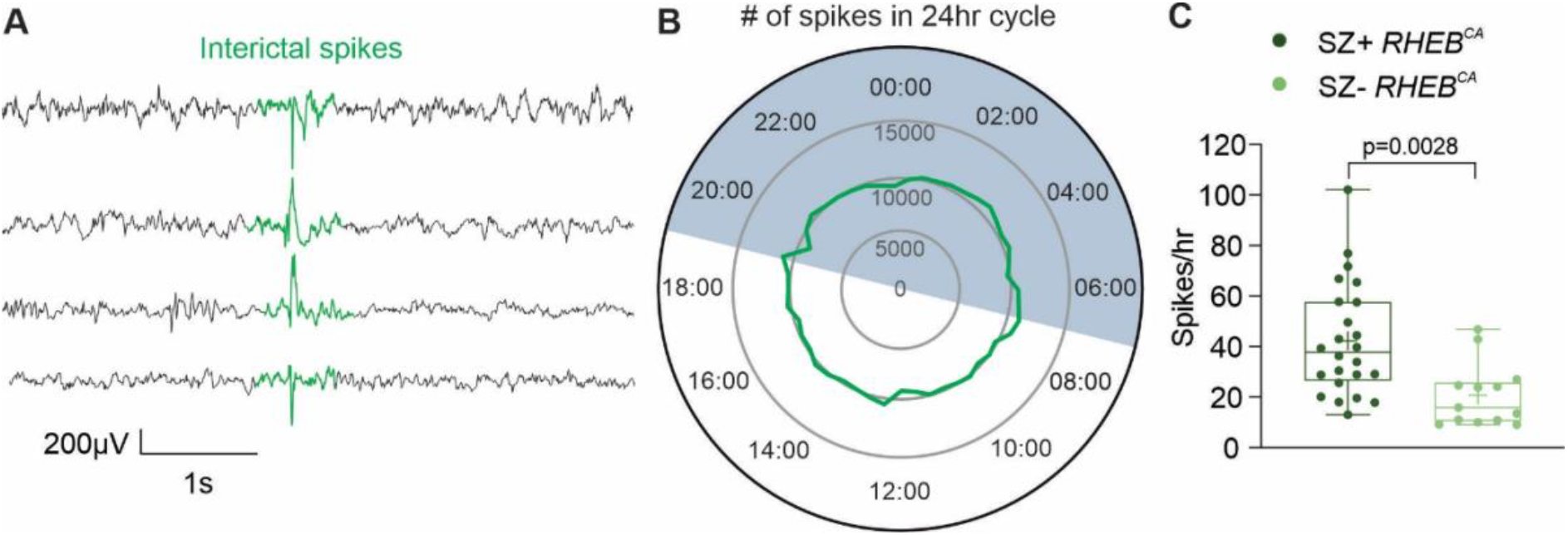
The *RHEB*^CA^ mouse model displays features of cortical hyperexcitability. **A.** Representative traces from 4 different animals showing interictal spikes (aligned at centre). **B.** Clock graph displaying the number of spikes recorded during the 12h:12h light-dark cycle; mice with and without generalised seizures (SZ+ *RHEB*^CA^ and SZ- *RHEB^CA^* respectively) were included in this analysis (n=38 mice). The dark phase was between 19:00 and 07:00. **C.** Box plot showing the number of interictal spikes per hour, (SZ- *Rheb*^CA^ n=13; SZ+ *Rheb*^CA^ n=24, unpaired Student’s t-test).

Behavioural and cognitive co-morbidities including learning disability and high autism spectrum quotient are common in FCD patients^22,23^. We therefore examined spontaneous alternation in a T-Maze (**Fig. 4A, B**) and performed tests of olfactory discrimination (**Fig. 4C, D**), both of which are reliant on prefrontal cortex function^24,25^, in SZ- *RHEB^CA^* and SZ+ *RHEB*^CA^ animals. Interestingly, only SZ+ *RHEB*^CA^ animals exhibited a significant impairment in alternation in the T-maze and in time spent investigating neutral odours and social olfactory cues **(Fig. 4B, D)**, consistent with learning disability and impaired social cognition often seen in patients with prefrontal FCD22,23.

**Figure 4.**
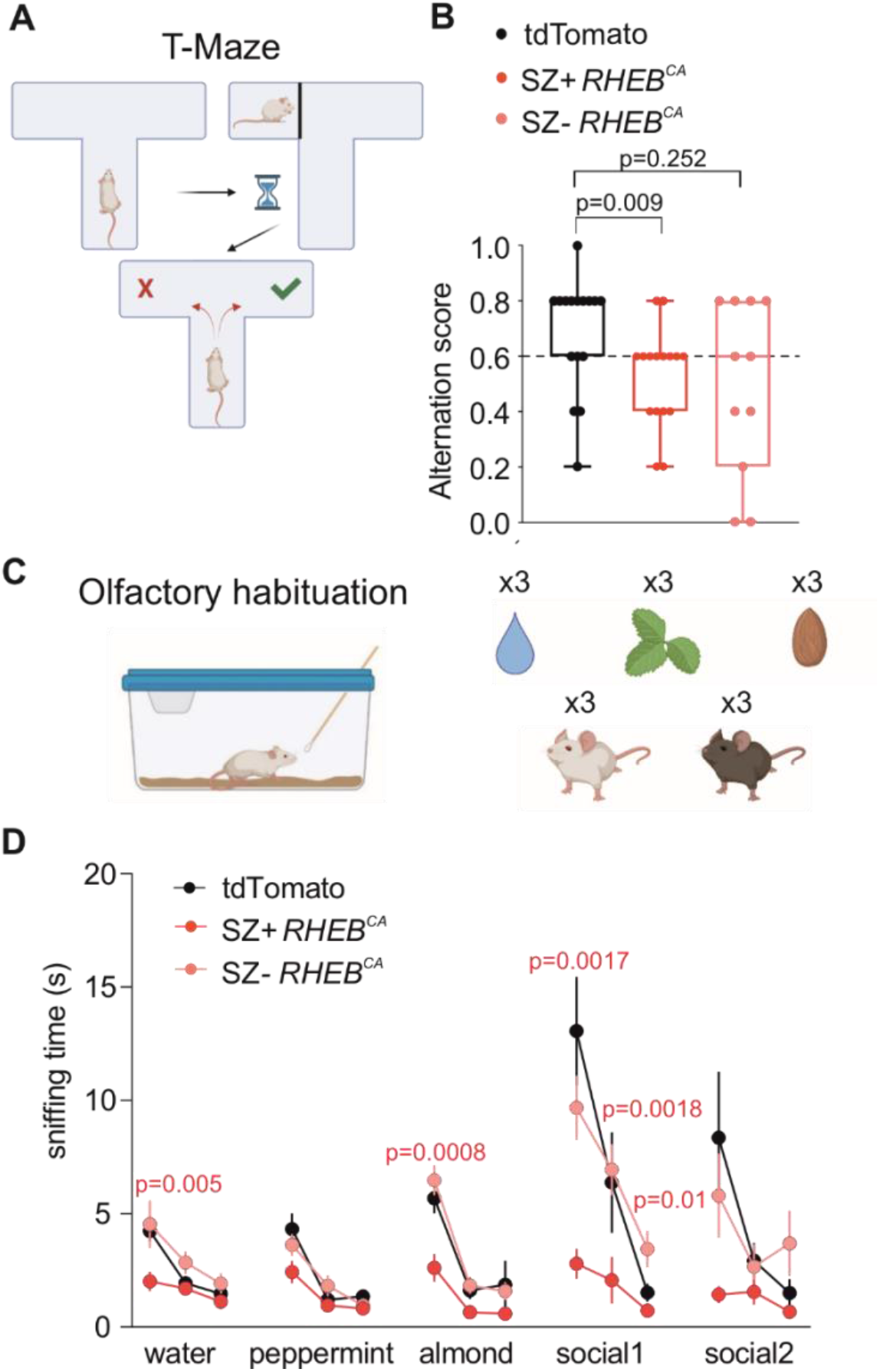
*RHEB*^CA^ mice have cognitive impairments. **A.** T-maze test. **B.** Box plot displaying the average alternation score in the T-Maze test for td-Tomato (n=16), SZ+ *RHEB^CA^* (n=16) and SZ- *RHEB*^CA^ (n=11) mice (Fisher’s Exact Test (>0.6 vs <=0.6), tdTomato vs SZ+ *RHEB*^CA^ or SZ- *RHEB*^CA^). **C.** Olfactory habituation test. Each smell (water, peppermint, almond and other mice [social]) was presented 3 times. **D.** Graph displaying the sniffing time of 3 neutral smells (water, peppermint, almond) and 2 social smells in tdTomato (n=16), SZ+ *RHEB^CA^* (n=17) and SZ- *RHEB^CA^* (n=11) mice (Repeated measured 2-way ANOVA followed by Bonferroni multiple comparison test).

### *CAMK2A-EKC* gene therapy reduces seizures without affecting cognition

The *RHEB*^CA^ model thus has both construct and face validity for FCD II. We asked whether expressing EKC in the dysplastic region to decrease neuronal and circuit excitability attenuates seizures. After recording 15 days of baseline ECoG (**Fig. 2**), we injected either AAV9-*CAMK2A*-EKC, tagged with green fluorescent protein (GFP), or a control virus, AAV9-*CAMK2A*-GFP, in the electroporated frontal lobe (**Fig. 5A**). Animals were randomized to the treatment groups and the experimenter was blinded to the allocation. ECoG recording was suspended for two weeks to preserve battery life and to allow for transgene expression, before resuming for another 15 days. Both SZ+ *RHEB^CA^* and SZ- *RHEB^CA^* animals were injected. In a subset of animals, we analysed the spatial expression of the reporter protein as well as pS6 expression. Overall, we observed expression of the viral reporter GFP in ~20% of the tdTomato-positive area, but also outside the dysplastic region, with considerable inter-animal differences (**Fig. S1A**). We observed no difference in pS6 expression between the *CAMK2A-EKC* and *CAMK2A*-GFP groups (**Fig. S1B**).

**Figure 5.**
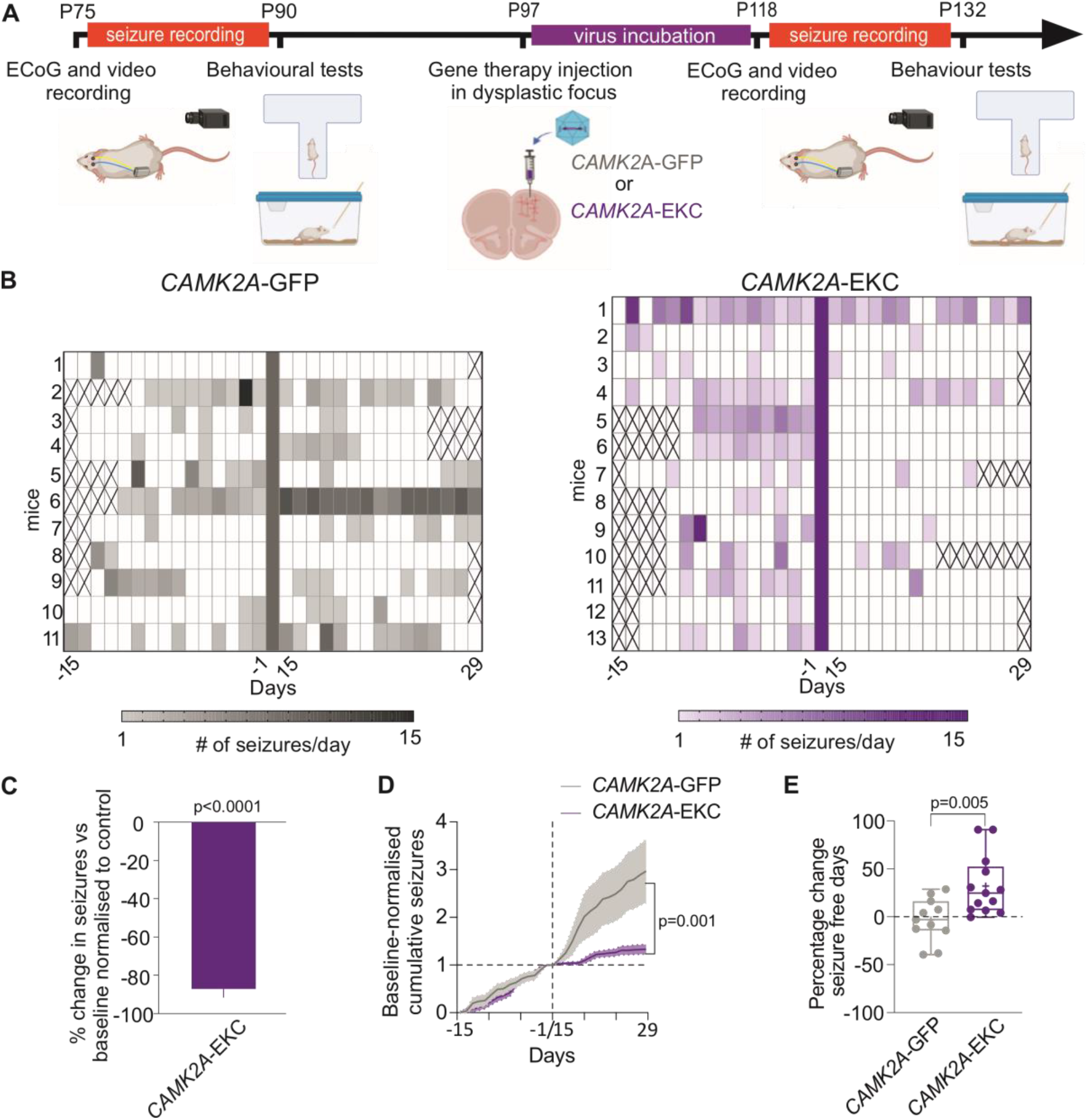
*CAMK2A-EKC* therapy reduces seizure frequency. **A.** Timeline. **B.** Heatmap displaying seizure occurrence in each animal injected with either *CAMK2A*-GFP (grey) or *CAMK2A*-EKC virus (purple) before and after the treatment (virus injection on day 0). Crosses correspond to days where data acquisition was incomplete. **C.** Graph displaying the change in number of seizures in animals treated with *CAMK2A*-EKC normalised to the average change in *CAMK2A*-GFP treated animals (one sample Wilcoxon test vs 0). **D.** Cumulative seizures normalised to baseline (*CAMK2A*-GFP n=11 mice, *CAMK2A*-EKC n=13 mice, repeated measures 2-way ANOVA followed by Bonferroni multiple comparison test). **E.** Graph showing the percentage of seizure-free days (unpaired Student’s t test).

Epileptic *RHEB^CA^* animals (SZ+ *RHEB^CA^*) injected with AAV9-*CAMK2A*-EKC showed a robust decrease in seizures in relation to their baseline (−64% ± 12%, n=13 mice, p=0.0002, one-sample paired t-test;) (**Fig. 5B**). In contrast, animals treated with the control virus AAV9-*CAMK2A*-GFP showed an increase in seizures, although this fell short of significance (**Fig. 5B;** +192% ± 71%, n=11 mice, p=0.22, one sample paired t-test). When this spontaneous evolution of seizure frequency was taken into account, we estimated that *CAMK2A-EKC* decreased seizures by ~87% (**Fig. 5C**).

Overall, 60% of EKC-treated SZ+ *RHEB*^CA^ mice became seizure-free by the last week of recording (**Fig. 5B**), and there was a 31% increase in seizure-free days (**Fig. 5E**). The effect of the treatment was also evident when plotting the cumulative evolution of seizures after treatment, normalised to each animal’s baseline (**Fig. 5D**). No effect on seizure duration was however seen (**Fig. S2**). The light-dark cycle pattern of seizure occurrence was maintained in *CAMK2A*-EKC treated animals although was not evident in *CAMK2A*-GFP-treated animals (**Fig. S3**).

We also assessed the effect of the treatment on interictal spikes. These continued to show no obvious diurnal variability in either treatment groups (**Fig. 6A**). There was however a significant decrease in the frequency of interictal spikes after treatment with *CAMK2A*-EKC in SZ+ *RHEB^CA^* mice, while SZ- *RHEB^CA^* animals showed no difference (**Fig. 6B**).

**Figure 6.**
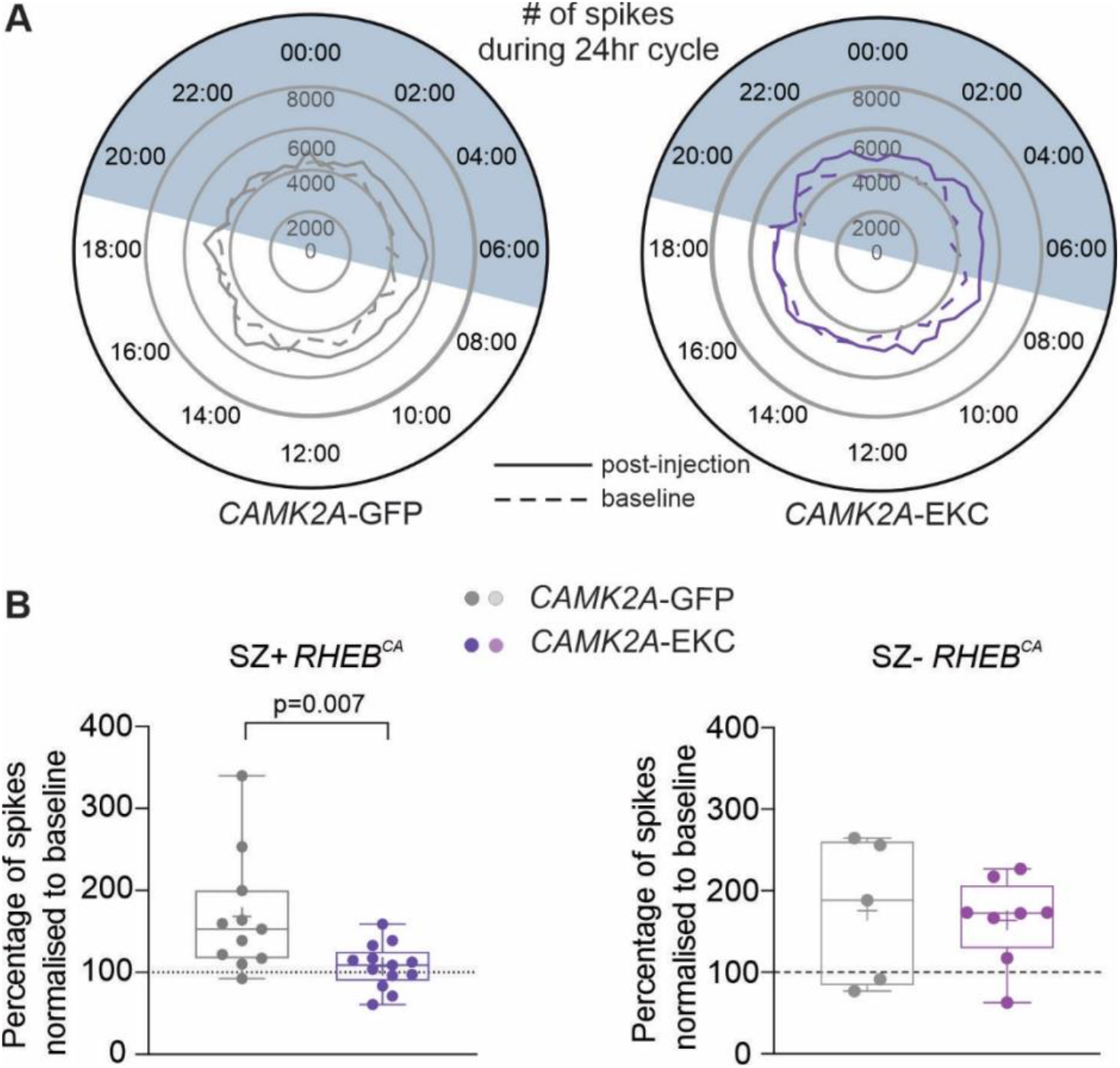
*CAMK2A-EKC* therapy reduces interictal spikes in *RHEB^CA^* animals with seizures. **A.** Circular graph displaying the number of spikes before (dash line) and after the therapy (continuous line) over 24h cycles. **B.** Box plot displaying average spike frequencies, normalised to baseline, in SZ+ *RHEB*^CA^ animals (*CAMK2A*-GFP n=11 mice, *CAMK2A*-EKC n=13 mice, unpaired Student’s t test) and in SZ- *RHEB*^CA^ animals (*CAMK2A*-GFP n=5 mice, *CAMK2A*-EKC, n=8 mice). We also asked whether the gene therapy either worsens or rescues the behavioural phenotype observed before treatment (**Fig. 4**). Neither the working memory test nor the social discrimination task was altered after treatment with *CAMK2A-EKC* or *CAMK2A*-GFP in animals with seizures **(Fig. 7A, B)**. Similarly, no differences in behaviour were seen in animals without seizures after treatment with *CAMK2A-EKC* or *CAMK2A*-GFP **Fig. 7A, C**).

**Figure 7.**
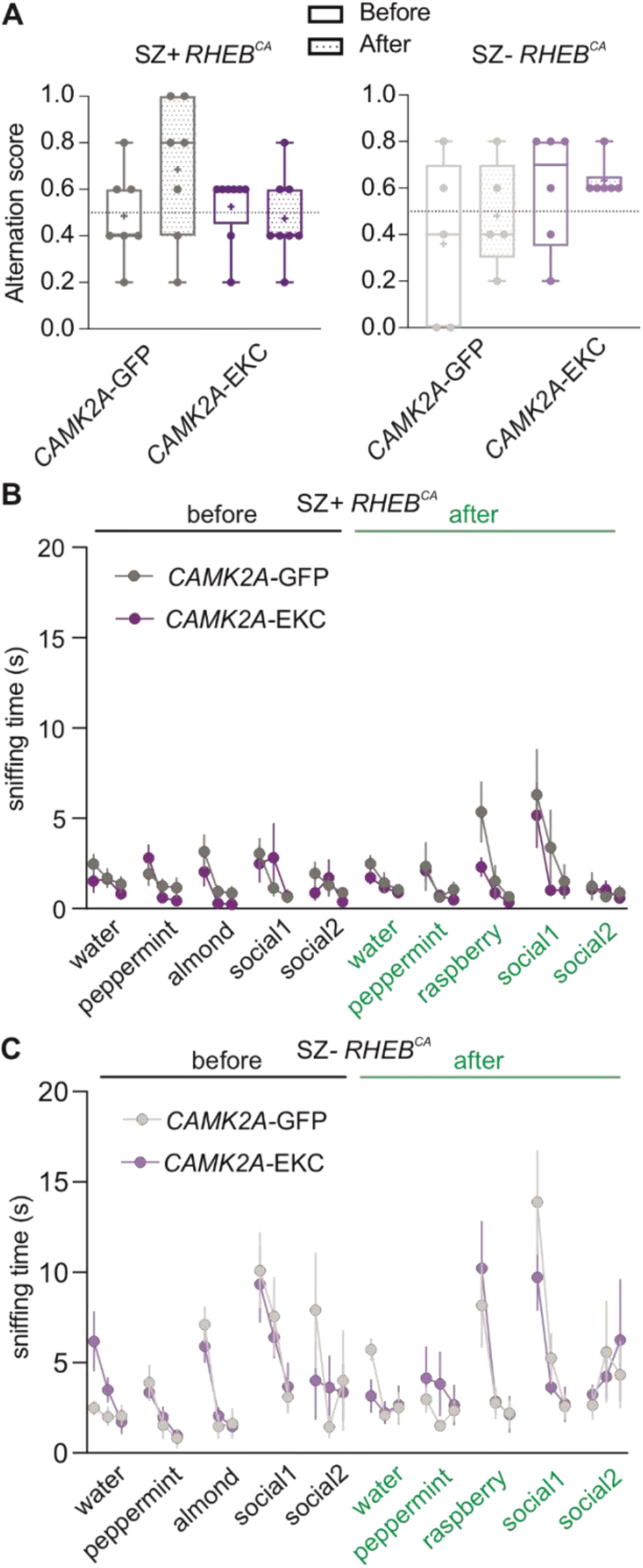
*CAMK2A-EKC* therapy neither aggravates nor worsens behavioural deficits in the *RHEB*^CA^ model. **A.** Box plot showing alternation scores in the T-Maze before and after *CAMK2A*-GFP (n=7) or *CAMK2A*-EKC (n=8) treatment in SZ+ *RHEB*^CA^, and *CAMK2A*-GFP (n=5) or *CAMK2A*-EKC (n=6) treatment in SZ- *RHEB*^CA^. Each circle represents an individual animal (p=0.1407 two-way ANOVA). **B.** Sniffing time for 3 neutral smells and 2 social smells before and after either *CAMK2A*-GFP (n=7) or *CAMK2A*-EKC (n=8) treatment (p=0.37 repeated measurement Mixed-effects analysis). **C.** Sniffing time for 3 neutral smells and 2 social smells before and after either *CAMK2A*-GFP (n=5) or *CAMK2A*-EKC (n=6) treatment (p=0.98 repeated measurement Mixed-effects analysis).

## Discussion

The main finding of the present study is that gene therapy with *CAMK2A-EKC* is highly effective in reducing spontaneous generalised seizures in the *RHEB*^CA^ model of FCD II, while neither worsening nor improving cognitive and behavioural comorbidity.

The study underlines the construct, face and predictive validity of the *RHEB*^CA^ model (**Figs. 1–3**)^9,26^. Dysregulation of the mTORC1 pathway leads to defects in migration of neural progenitors and cellular and circuit abnormalities ^1^. We extend the characterization of the *RHEB*^CA^ mouse model by showing that it has significant impairments in working memory and social recognition (**Fig. 3**). Although cognitive comorbidities accompany several forms of epilepsy and have been attributed to some extent to the acute effects of seizures and anti-seizure medication, they are especially common in ‘mTORopathies’ including FCD. In this study we aimed to reproduce as closely as possible a clinical scenario where treatment would be given following the emergence of seizures. The absence of significant changes in both behavioural tests is encouraging with respect to the safety of our therapy. On the other hand, failure to improve these aspects of the phenotype argues that they arise from the underlying circuit dysfunction rather than from seizures *per se*. Importantly, we showed that animals without spontaneous generalised seizures present no clear cognitive defects (**Fig. 3**), suggesting that although these impairments could be associated with the structural malformations and/or mTOR hyperactivity, seizures during development are a key factor for the manifestation of the behavioural phenotype. It remains to be determined whether an intervention earlier in life could prevent circuit dysfunction and thus mitigate behavioural deficits.

Although EKC treatment was highly effective, there was only relatively low spatial overlap between dysplastic neurons and virus expression. This may have reflected difficulty targeting the dysplastic zone reliably when guided only by tdTomato fluorescence. However, alternative possible explanations include limited tropism of the AAV9 capsid for dysplastic neurons, or limited ability of the *CAMK2A* promoter to drive expression of transgenes. Differences in AAV tropism towards neurons in mouse models have been previously reported ^27^ and neurons carrying the *RHEB* (S16H) mutation have altered membrane properties^28^. Importantly, this low overlap, in combination with the decrease of the number of generalised seizures observed in the *CAMK2A-EKC* group, suggests that treating the periphery of the FCD area is sufficient to prevent generalisation of seizures. However, we also found that animals exhibiting seizures generally exhibited a more profound decrease of the number of interictal spikes after EKC treatment, than animals without seizures. The effect of EKC therapy would therefore be consistent with a reduction in network excitability at the focus since interictal spikes are thought to be generated locally.

As mentioned above, anti-seizure treatment using EKC (or upregulating endogenous *Kcna1* transcription) has been previously reported in models of temporal lobe epilepsy^15,16^. The present study adds FCD II to the range of epilepsy models in which gene therapy with *CAMK2A-EKC* is effective. Although overactivation of the mTOR pathway is the commonest signalling disorder underlying FCD II, it is not the only one, and in clinical practice establishing the ultimate cause is only rarely possible without sampling brain tissue from the lesion. Nevertheless, the success of the *CAMK2A-EKC* reported here supports the potential use of the treatment in focal epilepsy irrespective of its aetiology. Promising results of ion channel-based gene therapy have previously been reported in the same model of FCD II^28^. A treatment aimed to counteract overexpression of the hyperpolarization-activated cyclic nucleotide-gated potassium channel isoform 4 (HCN4) was effective in reducing seizure occurrence^28^. Nevertheless, it remains to be determined whether this treatment’s effectiveness is limited to epilepsies caused by overactivation of mTORC1.

Interestingly, an increase in mTORC1 activity has been linked with alterations in Kv1.1 expression^29–32^. The interpretation of these studies is complex, as they suggest both an mTORC1-dependent and an independent regulation of Kv1.1 secondary to seizures, underlining the multifaceted nature of the mTOR pathway^33^. Indeed, an increase or a decrease in Kv1.1 hippocampal expression has been observed depending on the model used, and more work is needed to understand whether Kv1.1 expression is altered in FCD II associated with mTOR overactivation^29,31^.

Although invasive, injection of an AAV vector is likely to be a more palatable option than surgical resection. Indeed, the focal nature of FCD, which can frequently be detected on MRI, provides a target for AAV delivery. The local transduction of neurons avoids one of the main limitations of drug treatment, whether using antiseizure medication or mTOR-inhibiting drugs such as sirolimus (rapamycin) or its analogue everolimus, namely the risk of side effects arising from actions throughout the body^34^. In addition, whilst rapamycin and its analogues need to be taken continuously, gene therapy aims for a long-term solution in one intervention.

In conclusion, overexpression of the engineered potassium channel is an effective approach to treat seizures in FCD II. We provide a robust and safe treatment option, potentially suitable for clinical translation, for one for the commonest causes of refractory epilepsy.

## Acknowledgments

We are grateful Drs Laura Cancedda, Annalisa Savardi and Bruno Pinto from the Istituto Italiano di Tecnologia, Genoa, Italy, for the training on *in utero* electroporation. We would like also to thank Eugenia Almacellas for the analysis of the transduced/electroporated areas overlap and all DCEE for the useful discussions.

## Funding

GOSH/Spark Research Grant V4019 (DMK, GL and VM)

Medical Research Council Programme Grant MR/V034758/1 (DMK, GL and SS)

Epilepsy Research UK Emerging Leader Fellowship F1701 (GL)

Epilepsy Research UK Emerging Leader Fellowship F1901 (VM)

The Wellcome Trust Investigator Award 212285/Z/18/Z (DMK)

Medical Research Council New Investigator Project Grant MR/S011005/1 (GL)

## Materials and Methods

### Animals

All procedures were performed in accordance with the UK Home Office Animals (Scientific Procedures) Act 1986. All mice were housed under a non-reversed 12 h:12 h light–dark cycle and had access to food and water *ad libitum*.

### *In utero* electroporation

The surgery was performed as previously described^17^. Pregnant CD-1 dams (E14 ± 0.5) (Charles River) were anesthetized with isoflurane (induction 5%; maintenance 2.5%), and given Betamox (15 mg/ml), buprenorphine (0.03 mg/ml), Metacam (0.15 mg/ml) intramuscularly^17^. The dam was shaved, and an incision was performed close to the midline of the abdomen. The uterine horns were then exposed with a laparotomy. The CAG-tdtomato or CAG-tdtomato-T2A-*RHEB^CA^* plasmid (3.5μg/μl) was mixed with the dye Fast Green (0.3 mg/ml,Sigma-Aldrich) and injected (~1 μl/embryo) with a glass micropipette through the uterine wall unilaterally in the lateral ventricle. Then, 3 platinum electrodes soaked in sterile saline ~37°C were positioned around the embryo’s head (2 tweezer-type circular electrodes (cathodes) on the temporal lobes and a third additional electrode (anode) on the frontal lobe). The electroporation protocol consisted of one cycle of 6 electric pulses (28V) of 100ms duration with a 1s inter-pulse interval (electroporator from Nepagene NEPA21 type II). The uterine horns were continuously moistened with warm sterile saline during the electroporation. After all the embryos were electroporated, the uterine horns were inserted back into the abdominal cavity. The dam was sutured and given 0.5 ml saline subcutaneously and placed in a recovery chamber for 30 minutes before being returned to its home cage. The animal was closely monitored for 10 days afterwards.

Between P21 and P30, offspring were checked for tdTomato fluorescence with an epifluorescence stereomicroscope (Leica). The centre of the electroporated area was noted in reference to Bregma for further procedures and tdTomato-positive animals were single-housed. The overall success rate was 37%.

### Transmitter and cannula implantation

*RHEB^CA^* mice (weight 30-50 g) were anesthetised with isoflurane (induction 5%, surgery 2.5%) and placed into a stereotaxic frame (Kopf). They were injected subcutaneously with buprenorphine (0.03mg/ml) and Metacam (0.15 mg/ml). A wireless ECoG transmitter (catalog # A3028C-CC single-channel transmitter, Open Source Instruments) was implanted subcutaneously with a subdural intracranial recording electrode positioned above the central point of the electroporated area. A reference electrode was implanted over the visual cortex in the contralateral hemisphere. A cannula (Plastics One) was positioned above the injection site for delivery of AAV vectors later during the experiment. The electrodes and cannula were then secured with dental cement. The animal was given a subcutaneous injection of 0.5ml saline at the end of the surgical procedure and placed in a recovery chamber for 30 mins before being returned to its home cage. The animal was then monitored daily for 5 days.

### Viral vector injection

After 2 weeks of baseline ECoG recording, mice were briefly anaesthetised with 2.5% isoflurane, and injected with 900nl of adeno-associated virus serotype 9 expressing either *CAMK2A*-GFP or *CAMK2A*-EKC-GFP, via the cannula^14^. The injection was done at 3 different depths to cover the entire cortex thickness (z: −1, −0.5, −0.3 mm) at 100nL/min, with a 5 min pause between injections to avoid backflow. The animals were quarantined for 24h before being returned to their home cage for the next 2 weeks. The assignment of *CAMK2A*-GFP or *CAMK2A*-EKC-GFP to animals was randomised and the experimenter was blinded to the allocation.

### ECoG recording acquisition and analysis

ECoG was continuously recorded during 15 days before the viral vector injection, and then paused for 2 weeks before recording for a further 15 days. Data were acquired at a sampling rate of 256 Hz, and band-pass filtered between 1 and 160 Hz. To reduce electrical noise and the risk of headstage dislocation, animals were individually housed in a Faraday enclosure.

Epileptiform activity was identified and classified automatically using custom-written MATLAB code. Frequency and time-domain features (variance, extreme values, coastline, and kurtosis in the time domain, and alpha/beta power (8-30), low gamma (20-40Hz), high gamma (40-120Hz), spectral entropy) extracted at two timescales (1s and 10s) were used to calculate principal components, from which putative epileptiform activity was located and classified. Spikes not associated with continuous high-gamma band (40-100Hz) activity were considered isolated ‘interictal spikes’, while groups of spikes associated with continuous high-gamma activity were considered ‘polyspikes’. Seizures were considered to be any activity which continued for at least 10s, and which displayed seizure ‘anatomy’ (see Fig. 2D). Seizure anatomy was defined as the presence of three stereotyped patterns in visually confirmed seizures in the *RHEB*^CA^ model: a sentinel spike, a rising or falling gamma band oscillation indicative of the tonic phase, and terminal rhythmic after-discharges in the low frequency range (1-5Hz). ‘Generalised seizures’ were classified as activity which met the criteria for a seizure and showed increased signal amplitude in the tonic phase. These criteria reliably excluded intermittent periods of noise and electromyographic artefacts from analysis. Activity classified with a low confidence (only two of the three criteria for a seizure were met) was manually confirmed (blind to experimental group).

### Video recordings

IP cameras from Microseven (https://www.microseven.com/index.html) were used as previously described^16^.

### Behavioural assays

All behavioural tests were performed 1 hour after the start of the light phase and 1 hour before its end. When possible, the tests were performed under red light to reduce anxiety.

#### Spontaneous T-maze alternations

This test was performed as previously described under red light^35^. The apparatus made of transparent Plexiglass was cleaned with water after each animal. The walls of the apparatus were no less than 20 cm height, with a full length and width of 50 cm. Each of the arms was 25 cm long, and the running track was 50 cm long. The protocol for T-maze alternation was adapted from^35^. Mice were habituated to the room for 5-10 minutes before the experiment started. They were then placed at the starting area (bottom of the ‘T-shaped’ maze), facing away from the track. A central partition divided the end of the track to differentiate two directional choices. The animals initiated movement towards the apex of the T and chose one arm to enter. A positive entry was confirmed once the whole body (tail excluded) of the mice had entered the arm. The guillotine door was lowered to form an entrapped space for mice to stay for 30 seconds. Mice were then returned to the starting area and allowed to run down the track again with all doors open and without the separation wall in the middle. Their second positive arm entry was recorded as either the same or different from the first entry. The mice were then immediately returned to the starting area with no delay for the next trial. For each mouse, 5 successive trials were performed. If the animal suffered a seizure in the middle of the test, it was placed back in its homecage, and returned in the arena after all the other animals had been tested.

#### Olfactory discrimination

The olfactory discrimination test was adapted from^36^. The olfactory stimuli were prepared by diluting essential oils of peppermint, almond and raspberry in water (1:20). The solutions were freshly prepared and clean cotton swaps were briefly dipped in the different solutions. Both habituation and test were performed under white light. Mice were habituated for 30 minutes to an empty cage without bedding and a clean cotton swab inserted through the drinking bottle hole to remove any novelty confounder represented by the swab tip. The scented cotton swabs were then presented at the same height as the initial clean swab. Each smell was presented three times, lasting 2 minutes for each trial with an inter-trial interval of approximately 1 minute (the time it took for the experimenter to change the cotton swab). The order of the scents was water, peppermint, almond (neutral smells) and 2 smells from the bedding of other mice of the same sex (social smells). To control for the test repetition after the therapy injection, the almond odour was substituted with raspberry scent and the social smells were from different mice. The animals were expected to show habituation to each repeated odorant across the three trials and a novelty response when faced with a different smell. In addition, they were expected to show more interest to the social smells than the neutral smells. The odour discrimination, habituation and interest were assessed by quantifying the time spent sniffing the cotton swab. The amount of time spent sniffing was taken on-site by the experimenter. The experimenter was blinded to the treatment groups.

### Immunofluorescence staining

#### Tissue collection

The animals were exsanguinated via intracardiac perfusion with Phosphate Buffered Saline (PBS) followed by paraformaldehyde (PFA 4%). The brain tissue was kept overnight in PFA 4% after dissection, then washed with PBS before slicing. Coronal sections of 70μm thickness from PFA-fixed mice brains were performed using a vibrating microtome (Leica VT1000S) and stored in PBS with (0.02% sodium azide (NaN3) at 4°C.

#### Immunostaining and Image Acquisition

Free-floating brain sections were first washed 3 times for 5 minutes with PBS, followed by 3 hours of incubation in blocking buffer (0.3% triton-100X, 3% BSA, and 5% goat serum). Slices were then incubated overnight at 4°C with a primary antibody in diluted blocking buffer (1 in 4) (dilution 1:1000, anti-rabbit pS6 (#35708, Cell Signalling Technologies)). Slices were then rinsed 3 times in PBS followed by secondary antibody incubation with Alexa Fluor 488 goat anti-rabbit (1:1000) diluted in PBS for 3 hours at room temperature in the dark on a shaker. Slices were mounted and preserved using Vectashield antifade mounting medium with DAPI (Vector Laboratories). Tile-scan images were acquired with Zen software (Zeiss) on an LSM710 confocal microscope (Zeiss). Quantification of pS6 fluorescence intensity was done using FIJI software, the region of interest (ROI) was determined with a mask thresholding for the particles corresponding to neurons. The total intensity density in the ROI was normalised to the number of particles.

### Statistical analysis

Statistical analysis was performed on raw and normalised data with Prism (GraphPad Software, Inc., CA). Data are plotted as box and whiskers, representing interquartile range (box), median (horizontal line), the mean (+ symbol) and maximum and minimum (whiskers), together with all the points. The statistical analysis performed is shown in each figure legend. Deviation from normal distributions was assessed using D’Agostino-Pearson’s test, and the F-test was used to compare variances between two sample groups. Student’s two-tailed t-test (parametric) or Wilcoxon test (non-parametric) was used as appropriate to compare means. To compare two groups at different time points, we used two-way repeated measure ANOVA or mixed-effects analysis, followed by Bonferroni post hoc test for functional analysis.

## Supplementary figures

**Fig.S1.**
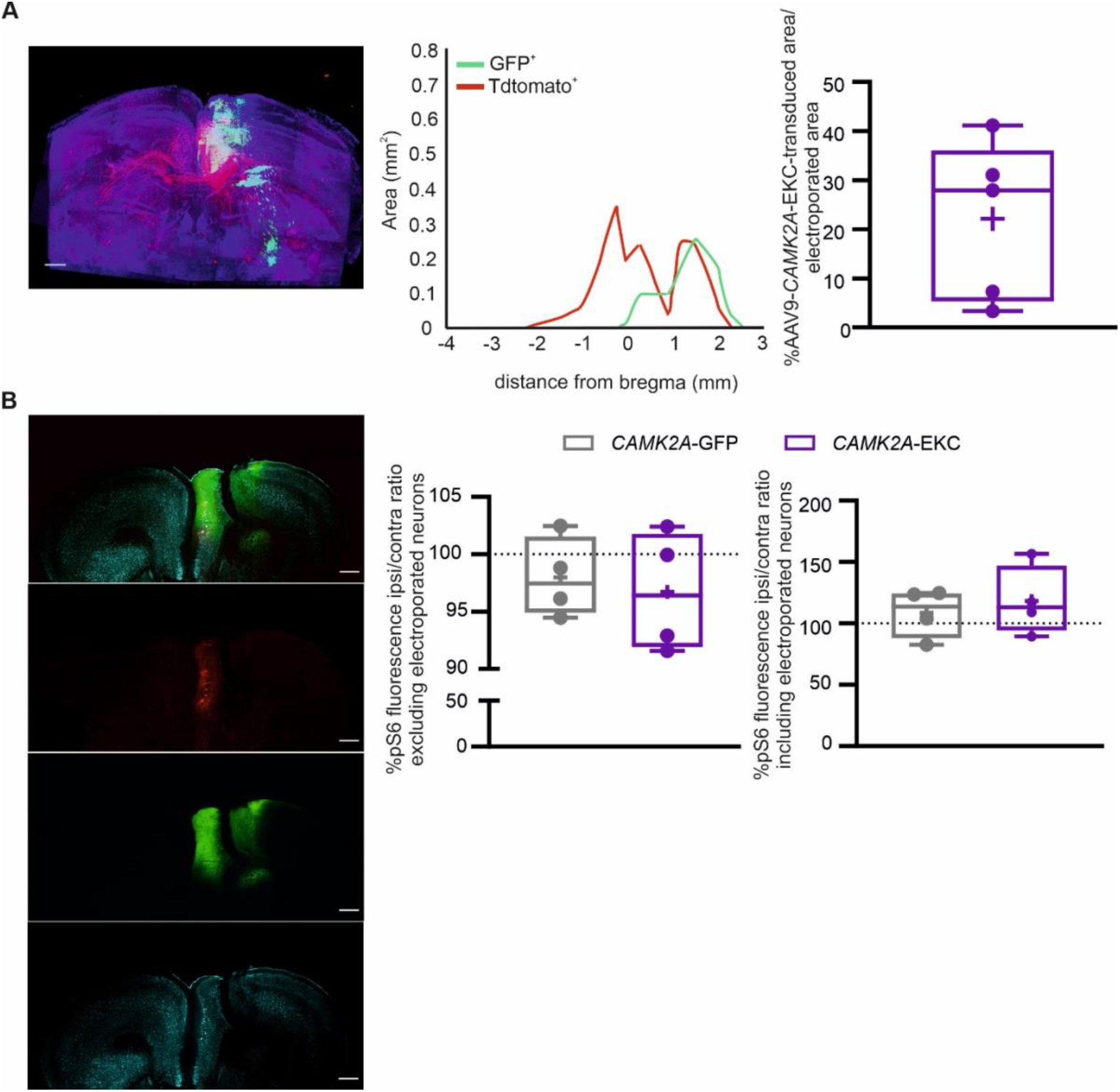
*CAMK2A-EKC* therapy spreads in the dysplastic region and does not reduce mTORC1 activity. **A.** 3D reconstruction of the spread of the dysplastic area (tdTomato, red fluorescence) and its overlap with expression of the GFP reporter included in the *CAMK2A*-EKC therapy, scale bar=500μm. (*middle*) Line graph displaying the anatomical overlap of the transduced (GFP^+^) and electroporated (tdTomato^+^) areas. (*left*) Box plot showing the percentage of electroporated area (tdTomato) overlapping with *CAMK2A*-EKC transduced area (GFP+) (n=5 *CAMK2A*-EKC-GFP mice). **B.** Triple immunofluorescence picture of prefontal cortical slice presenting a cortical dysplastic area (top panel: merged channels, red fluorescence tdTomato^+^ neurons, green fluorescence GFP+ neurons, cyan fluorescence pS6 protein expression). (*right*) Box plots displaying the % of pS6 fluorescence between the two hemispheres (ipsi = electroporated hemisphere) in animals from both groups (grey = *CAMK2A*-GFP, purple = *CAMK2A*-EKC, n=4 mice per group).

**Fig.S2.**
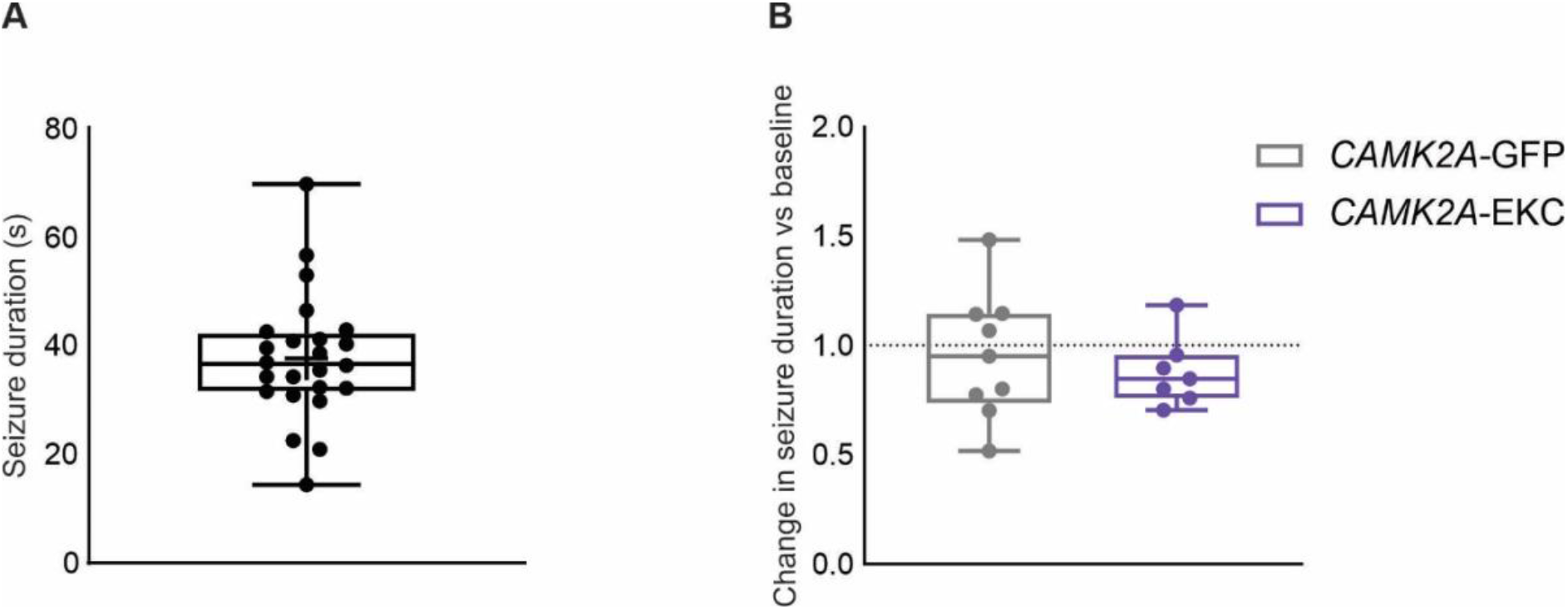
*CAMK2A-EKC* therapy does not change seizure duration. **A.** Box plot displaying the average durations of seizures recorded prior to AAV injection (n=24 mice). **B.** Box plot displaying the change in seizure duration normalised to baseline for *CAMK2A-EKC* (n=7 mice) and *CAMK2A*-GFP (n=9 mice) groups (p=0.5455, unpaired t-test).

**Fig.S3.**
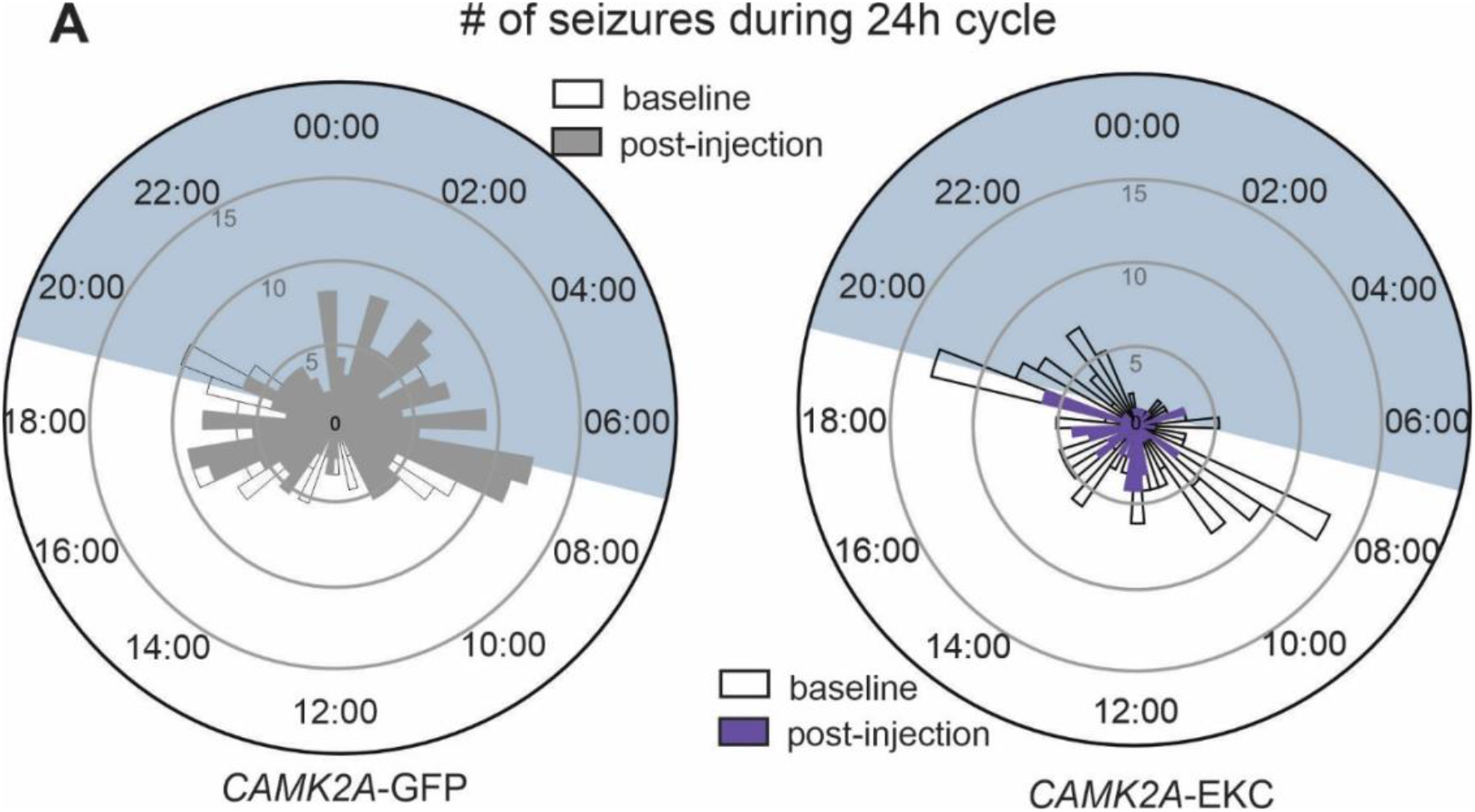
*CAMK2A-EKC* therapy does not change the light-dark cycle pattern of seizure occurrence. Circular graph displaying the number of seizures before (empty rectangles) and after the therapy (filled rectangles) over 24h cycles for animals injected with either *CAMK2A*-GFP (n=11 mice) or *CAMK2A*-EKC (n=13 mice).

## Notes

### Competing Interest Statement

The authors have declared no competing interest.

## References

1. Lee, W. S. et al. Cortical Dysplasia and the mTOR Pathway: How the Study of Human Brain Tissue Has Led to Insights into Epileptogenesis. Int. J. Mol. Sci. 23, (2022).

2. Blumcke, I. et al. Histopathological Findings in Brain Tissue Obtained during Epilepsy Surgery. N. Engl. J. Med. 377, 1648–1656 (2017).

3. Stevelink, R. et al. Epilepsy surgery in patients with genetic refractory epilepsy: A systematic review. Eur. J. Paediatr. Neurol. 21, e17 (2017).

4. Najm, I. et al. The ILAE consensus classification of focal cortical dysplasia: An update proposed by an ad hoc task force of the ILAE diagnostic methods commission |. Epilepsia 00, (2022).

5. Hu, S. et al. Somatic Depdc5 deletion recapitulates electroclinical features of human focal cortical dysplasia type IIA. Ann. Neurol. 84, 140–146 (2018).

6. White, A. R. et al. PI3K isoform-selective inhibition in neuron-specific PTEN-deficient mice rescues molecular defects and reduces epilepsy-associated phenotypes. Neurobiol. Dis. 144, 105026 (2020).

7. Lim, J. S. et al. Somatic Mutations in TSC1 and TSC2 Cause Focal Cortical Dysplasia. Am. J. Hum. Genet. 100, 454–472 (2017).

8. Baek, S. T. et al. An AKT3-FOXG1-reelin network underlies defective migration in human focal malformations of cortical development. Nat. Med. 21, 1445–1454 (2015).

9. Hsieh, L. S. et al. Convulsive seizures from experimental focal cortical dysplasia occur independently of cell misplacement. Nat. Commun. 7, 11753 (2016).

10. Proietti Onori, M. et al. RHEB/mTOR hyperactivity causes cortical malformations and epileptic seizures through increased axonal connectivity. PLOS Biol. 19, e3001279 (2021).

11. Kim, J. K. et al. Brain somatic mutations in MTOR reveal translational dysregulations underlying intractable focal epilepsy. J. Clin. Invest. 129, 4207–4223 (2019).

12. Heeroma, J. H. et al. Episodic ataxia type 1 mutations differentially affect neuronal excitability and transmitter release. DMM Dis. Models Mech. 2, 612–619 (2009).

13. Wykes, R. C. et al. Optogenetic and potassium channel gene therapy in a rodent model of focal neocortical epilepsy. Sci. Transl. Med. 4, 161–152 (2012).

14. Snowball, A. et al. Epilepsy Gene Therapy Using an Engineered Potassium Channel. J. Neurosci. 39, 3159–3169 (2019).

15. Qiu, Y. et al. On-demand cell-autonomous gene therapy for brain circuit disorders. Science 378, 523–532 (2022).

16. Colasante, G. et al. In vivo CRISPRa decreases seizures and rescues cognitive deficits in a rodent model of epilepsy. Brain J. Neurol. 143, 891–905 (2020).

17. Szczurkowska, J. et al. Targeted in vivo genetic manipulation of the mouse or rat brain by in utero electroporation with a triple-electrode probe. Nat. Protoc. 11, (2015).

18. Nguyen, L. H. et al. mTOR hyperactivity levels influence the severity of epilepsy and associated neuropathology in an experimental model of tuberous sclerosis complex and focal cortical dysplasia. J. Neurosci. 39, 2762–2773 (2019).

19. Liu, Z. et al. Systematic comparison of 2A peptides for cloning multi-genes in a polycistronic vector. Sci. Rep. 7, 2193 (2017).

20. Karoly, P. J. et al. Cycles in epilepsy. Nat. Rev. Neurol. 17, 267–284 (2021).

21. de Curtis, M. & Avanzini, G. Interictal spikes in focal epileptogenesis. Prog. Neurobiol. 63, 541–567 (2001).

22. Veersema, T. J. et al. Cognitive functioning after epilepsy surgery in children with mild malformation of cortical development and focal cortical dysplasia. Epilepsy Behav. 94, 209–215 (2019).

23. Fujimoto, A. et al. Epilepsy in patients with focal cortical dysplasia may be associated with autism spectrum disorder. Epilepsy Behav. 120, 107990 (2021).

24. Naik, A. A.et al. Extrahippocampal seizure and memory circuits overlap. eNeuro 9, (2022).

25. Boyd, A. M.et al. Broadcasting of Cortical Activity to the Olfactory Bulb. Cell Rep. 10, 1032–1039 (2015).

26. Zhao, S. et al. A brain somatic RHEB doublet mutation causes focal cortical dysplasia type II. Exp. Mol. Med. 51, (2019).

27. Chen, Y. H. et al. Sialic acid deposition impairs the utility of AAV9, but not peptide-modified AAVs for brain gene therapy in a mouse model of lysosomal storage disease. Mol. Ther. J. Am. Soc. Gene Ther. 20, 1393–1399 (2012).

28. Hsieh, L. S. et al. Ectopic HCN4 expression drives mTOR-dependent epilepsy in mice. Sci. Transl. Med. 12, (2020).

29. Raab-Graham, K. F. et al. Y. Activity-and mTOR-Dependent Suppression of Kv1.1 Channel mRNA Translation in Dendrites. Science 314, 144–148 (2006).

30. Sosanya, N. M. et al. Degradation of high affinity HuD targets releases Kv1.1 mRNA from miR-129 repression by mTORC1. J. Cell Biol. 202, 53–69 (2013).

31. Nguyen, L. H. & Anderson, A. E. MTOR-dependent alterations of Kv1.1 subunit expression in the neuronal subset-specific Pten knockout mouse model of cortical dysplasia with epilepsy. Sci. Rep. 8, 1–12 (2018).

32. Sosanya, N. M. et al. Rapamycin reveals an mTOR-independent repression of Kv1.1 expression during epileptogenesis. Neurobiol. Dis. 73, 96–105 (2015).

33. Laplante, M. & Sabatini, D. M. Regulation of mTORC1 and its impact on gene expression at a glance. J. Cell Sci. 126, 1713–1719 (2013).

34. Franz, N. et al. Adjunctive-everolimus-therapy-for tuberous-sclerosis-complex associated refractory-seizures: -Results-from-the-post-extension-phase-of-EXIST--3. Epilepsia 62, 3029–3041 (2021).

35. Deacon, R. M. J. & Rawlins, J. N. P. T-maze alternation in the rodent. Nat. Protoc. 1, 7–12 (2006).

36. Yang, M. & Crawley, J. N. Simple behavioral assessment of mouse olfaction. Curr. Protoc. Neurosci. Chapter 8, Unit 8.24 (2009).

